# Impact of trimethoprim on the river microbiome and antimicrobial resistance

**DOI:** 10.1101/2020.06.05.133348

**Authors:** J. Delaney, S. Raguideau, J. Holden, L. Zhang, H.J. Tipper, G.L. Hill, U. Klümper, T. Zhang, C.J. Cha, K. Lee, R. James, E. Travis, M.J. Bowes, P.M. Hawkey, H.S. Lindstrom, C. Tang, W.H. Gaze, A. Mead, C. Quince, A. Singer, E.M.H. Wellington

## Abstract

Recent evidence suggests that anthropogenic activity can increase the levels of antimicrobial resistance (AMR) in the environment. Rivers and waterways are significant examples of environmental settings that have become repositories of antibiotics and antibiotic resistance genes (ARGs). Our recent study quantified drug concentrations in freshwater samples taken at a range of sites located on the Thames catchment; the highest levels of antibiotics and other drugs were recorded downstream of waste water treatment plants (WWTPs). One specific antibiotic: Trimethoprim (TMP) was shown at elevated concentrations reaching 2000ng/L at particular sites. We have also shown a correlative relationship between the residue of TMP and the prevalence of sulfonamide antibiotic resistance genes such as sul1. Despite this, there is still no evidence of a causative relationship between TMP concentrations and the prevalence of the ARGs at river sites. The aim of the current study was to conduct in-depth analysis using a combination of large metagenomic, geospatial and chemical datasets, in order to conduct a comparison between those sites with the highest TMP and lowest TMP levels across the Thames catchment. We aimed to establish the proximity of these sites to WWTPs, their population equivalence (PE) and land coverage. A secondary aim was to investigate seasonal variation in TMP and ARGs. Exploring these factors will help to decipher the clinical relevance of ARG accumulation at river sites. A significant correlation was shown between TMP levels at river sites and their distance downstream from a WWTP. Three sites located on the Rivers Cut and Ray showed significantly higher TMP concentrations in winter compared to summer. The population equivalence (PE) for sites with the highest TMP levels was significantly higher than those with the lowest levels. The land coverage of sites with the highest TMP levels was significantly more urban/suburban than sites with the lowest TMP concentrations, which were found to be significantly more arable. Five ARGs relevant to TMP and sulfonamides were identified across the Thames catchment. The most prevalent ARG was sul1, which was significantly more prevalent in winter compared to summer. By contrast sul2 was found to be significantly more prevalent in summer compared to winter at a site on the River Coln. The prevalence of the class 1 integron marker gene (inti1) did not differ significantly by season or between sites with the highest/lowest TMP levels.

## Introduction

Antimicrobial resistance (AMR) is a global health issue, with forecasts predicting ten million deaths a year by 2050 if resistance levels continue to rise (O’Neill and The Review on Antimicrobial Resistance, 2016). The struggle against AMR has conventionally been viewed as a medical problem (Bengtsson-Palme et al, 2018). However, the environment is now being acknowledged as a substantial contributor to the global spread of antibiotic resistance (Wellington et al, 2013 & Singer et al, 2016;2019). Rivers and waterways are prominent examples of environmental settings that have been identified as a major repository and route for dissemination of both low levels of antibiotics and antibiotic resistance genes (ARGs) (Pärnänen et al, 2019; Amos et al, 2015).

A recent study quantified antibiotic concentrations in freshwater samples taken at a range of sites located on the Thames river catchment; the highest levels of antibiotics (as well as many other drugs) were recorded downstream of waste-water treatment plants (WWTPs) (Holden et al, 2020). While the origin and fate of these antibiotics could be strongly linked to community and clinical use, the impact that they have on the emergence of antimicrobial resistance in the environment remains unclear (Aslam et al, 2018). It has been proposed that ARGs have emerged within environmental bacteria to provide a protective effect against these polluting drugs (Märtinez et al, 2012). This is of concern to human health, as the genes under selection may be clinically relevant and have the potential to be transferred between environmental and clinical bacteria via horizontal gene transfer (HGT) (Baquero et al, 2008). However, it is also widely recognised that the evolutionary arms race, or red queen hypothesis: the concept that species need to continually adapt, evolve, and proliferate to survive plays an important role in shaping the environmental resistome (Papot et al, 2017). The fact that ARGs have continually emerged within the riverine environment to protect against antibiotic producing organisms in the riverine environment supports this latter concept (Singer et al, 2016). As a result, the primary drivers of AMR within complex riverine bacterial communities remain difficult to define.

We have previously shown that the prevalence of an integron borne ARG, *sul1*, correlates with the concentration of several antibiotics across river sites on the Thames catchment (Holden et al, 2020). *sul1* provides resistance to sulfonamides, a bacteriostatic class of antibiotics that were originally synthesised in the 1930s. Sulfonamides bind to dihydropteroate synthase (DHPS) in bacteria and hinder dihydrofolic acid formation which is vital for vitamin B formation and bacterial growth (Sköld, 2000). One sulfonamide; sulfamethoxazole (SMX) can be prescribed with trimethoprim (TMP) as co-trimoxazole to treat chronic bronchitis, shigellosis, and *Pneumocystis carinii* pneumonitis (Alsaad et al, 2013). SMX and TMP act in a synergistic manner to target successive steps of the folate synthesis pathway, allowing co-trimoxazole to be effective against most aerobic gram-negative and gram-positive organisms (Macejko and Schaeffer, 2008) (Figure 1).

**Figure 1.**
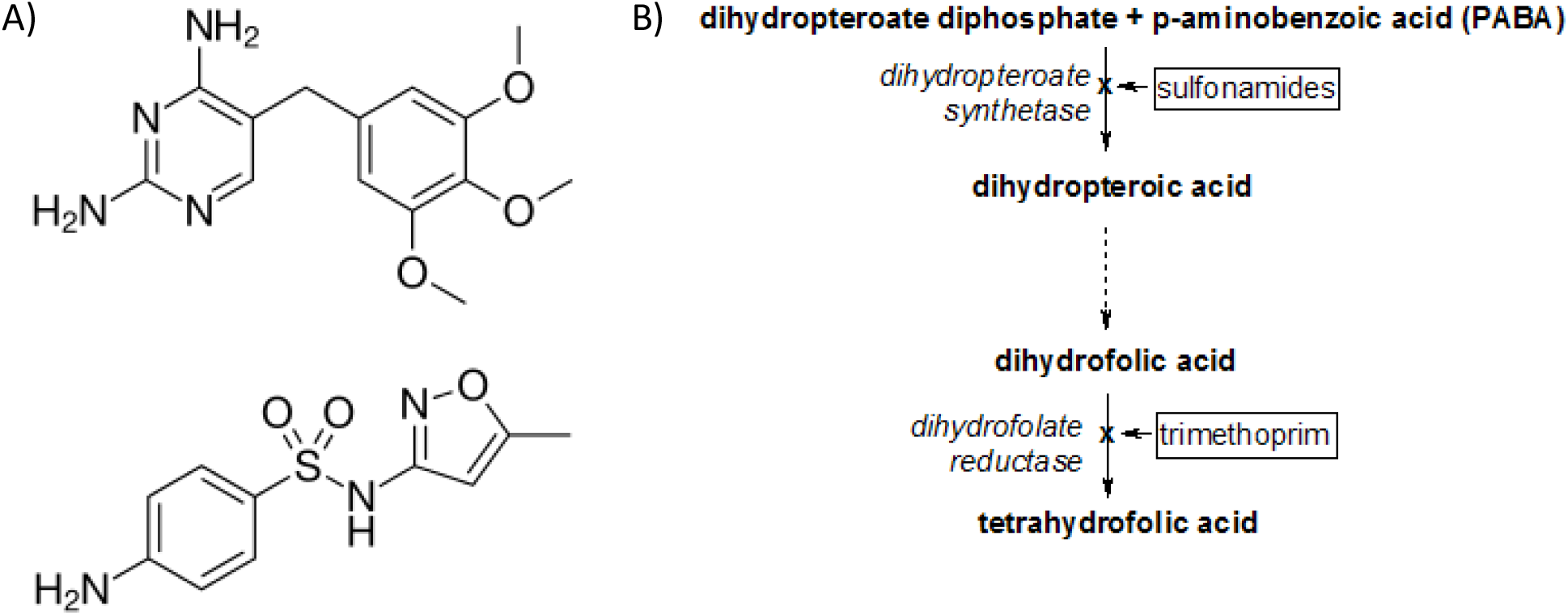
A) Chemical structure of SMX/TMP
B) SMX and TMP combined mechanism of action

Despite the synergistic effects of SMX and TMP, recent human antibiotic prescription surveillance data suggests that TMP is more commonly prescribed as a single-dose antibiotic for clinical purposes (Ashiru-Oredope et al, 2014). Furthermore, the recent veterinary medicines directorates report on antibiotics showed that a combined volume of 66.2mg/kg of sulfonamides/TMP was used to treat infections in pigs in 2015, which accounted for the highest proportion of all antibiotics prescribed for pigs (24%) (Bos et al, 2019). Studies have identified several different ARGs relevant to TMP (*DfrA1*, *DfrE*, *DfrB2/3, DfrB1/6)* (Šeputienė et al, 2010) and sulfonamides (*sul1*, *sul2, sul3, sul4*) (Razavi et al, 2017). Due to the prominence of sulfonamide resistance in the environment, the *sul1* gene has become incorporated into the structure of many class 1 integrons (CL1s) (Na et al, 2014). Several studies have noted the persistence of sulfonamide and TMP resistance genes in the same environmental settings, often located on the same plasmids (Phuong Hoa et al, 2008; Kiiru et al, 2013; Muziasari et al, 2014).

During our previous work, sites on the Thames catchment with elevated TMP levels were identified (Holden et al, 2020). Despite this, there is still no evidence for a causative relationship between the residue of antibiotics such as TMP and the prevalence of ARGs (Holden et al, 2020). It is possible that the correspondence between ARGs and drugs downstream of WWTPs is purely correlative, as both are being released from the treatment process at the same time. In fact, studies have suggested that WWTPs themselves, as opposed to river sites are ‘hotspots’ of ARGs and therefore most of the selection for resistance occurs before antibiotic residue enters the riverine environment (Lood et al, 2017; Pazda et al, 2019; Fouz et al, 2020).

The aim of the current study was to analyse a combination of large metagenomic, geospatial and chemical datasets, to compare twenty sites across the Thames river catchment. Specifically, we aimed to look at sites with the highest and lowest TMP levels and establish the role that factors such as proximity to WWTPs, population equivalence (PE) and land coverage play in the accumulation of TMP at river sites. A secondary aim was to investigate seasonal variation in the levels TMP and SMX at river sites. Lastly, as TMP can be prescribed alongside SMX as co-trimoxazole, we aimed to establish if the levels of TMP and SMX identified at these specific sites mirrored the prevalence of ARGs relevant to TMP (*Dfr* genes) and SMX (*sul* genes).

## Methods

A total of 69 sampling locations were chosen within the Thames catchment. However, metagenomic data was only generated for twenty of these 69 sites, and these twenty sites were analysed during this study. All locations provide a depth of understanding regarding long-term water quality parameters. A subset of these locations also contributed to a previous effort on modelling AMR within the Thames Catchment (Amos et al 2015). The remaining sampling locations were selected to maximise the diversity of river environments, river inputs, and adjacent land uses. Specific sampling locations within a river catchment were selected to capture upstream and downstream positions from a known river discharge or confluence. All locations also had to consider site accessibility as a factor. Key discharges and confluences which structured sampling location choice were wastewater treatment plants, agricultural fields (e.g., animal grazing), urban runoff, canals, fish farms, and other rivers (see Table 1 Figure 1).

**Table 1.** Summary of the twenty sampling sites analysed in this study

| Name of Site | Thames<br>Tributary | Nearest<br>upstream<br>Thames WWTP<br>and treatment<br>type (Secondary<br>(S) or Tertiary (T)) | XY<br>coordinates | Postcode |
| --- | --- | --- | --- | --- |
| 1. Coln1 | River Coln | Withington (S) | X:410239<br>Y:207637 | GL7 5NY |
| 2. Coln2 | River Coln | Withington (S) | X:411501<br>Y:206777 | GL7 5NW |
| 3. Coln3 | River Coln | Bibury (S) | X:414943<br>Y:201324 | GL7 4AF |
| 4. Coln4 | River Coln | Fairford (S) | X:416285<br>Y:200409 | GL7 4LX |
| 5. Cut1 | River Cut | N/A | X:488447<br>Y:171940 | BG42 6JQ |
| 6. Cut2 | River Cut | Ascot (S) | X:485223<br>Y:171833 | RG42 5NR |
| 7. Cut4 | River Cut | Bracknell (T) | X:485544<br>Y:173282 | RG42 |
| 8. Cut6 | River Cut | Maidenhead (T) | X:490245<br>Y:179218 | SL6 2AR |
| 9. Leach1 | River Leach | Northleach (S) | X:420266<br>Y:204232 | GL7 3NS |
| 10. Leach2 | River Leach | Lechlade (S) | X:422552<br>Y:199090 | GL7 3HA |
| 11. Ray1 | River Ray<br>(Thames headwater) | N/A | X:410831<br>Y:187351 | N/A |
| 12. Ray3 | River Ray | N/A | X:412867<br>Y:185375 | SN5 7UJ |
| 13. Ray4 | River Ray | Swindon (T) | X:412403<br>Y:186741 | SN2 |
| 14. Ray5 | River Ray | N/A | X:412310<br>Y:187249 | SN2 |
| 15. Ray6 | River Ray | Swindon (T) | X:411910<br>Y:192617 | SN6 6 |
| 16. RiverThames1 | River Thames | Castle Eaton (S) | X:417451<br>Y:196098 | SN6 7RX |
| 17. RiverThames3 | River Thames | Witney (T) | X:444319<br>Y:208614 | OX29 4BT |
| 18. Thame1 | River Thame | Rowsham (S) | X:479984<br>Y:215448 | HP19 9UA |
| 19. Thame2 | River Thame | Aylesbury (T) | X:477886<br>Y:213623 | N/A |
| 20. Thame7 | River Thame | Little Milton (S) | X:460064<br>Y:197680 | OX10 7AW |

In the specific case of WWTPs, small WWTP were plants serving a population less than 10,000 and any plant serving over 10,000 was considered large. WWTP impact was determined based on size of the nearest WWTP and distance from the plant. Impact was determined based on distance away from the WWTP.

### Sampling regime

Samples were collected on two- or three-day sampling campaigns in August 2015, early March 2016 and August 2016. Sampling was carried out using purpose-made expandable sampling poles to allow access across the riverbed. The poles were fitted with a machined holder that secured a 50ml falcon tube. Sediment samples were collected by extending the pole into the river and scraping the upper sediment layer into the tube until it was largely full. Sediment tubes were stored on ice. Samples were collected at each site in triplicate. Water samples were collected in 500 ml screw cap bottles and stored on ice. Samples for nutrient analysis was prepared as previously described (Amos et al, 2015).

### Sample Processing

DNA extraction was carried out using the FastDNA™ spin kit for soil following manufacturers protocol, 0.5 g of sediment (wet weight) was used. DNA was pooled after extraction in 75μl to give total DNA in final volume of 150 μl.

### Metagenome

PCR free shotgun libraries were prepared for each sample from 2.5 ug metagenomic DNAs by Exeter Sequencing Service for the metagenomic sequencing using the TruSeq DNA library Prep Kit (Illumina). The libraries were sequenced to approximately 7GBp using the HiSeq 2500 rapid run mode (2X 250 bp paired end) by the Exeter Sequencing Service (http://sequencing.exeter.ac.uk/). A repeat sequencing effort was done on six DNA samples to determine if the coverage was sufficient and each library was sequenced to provide 35 GBp per sample and the total was assembled as below.

### Metagenome assembly, annotation and coverage

Metagenome assembly was performed using megahit v1.1.3 with option -k--min 29 (Dinghua et al, 2015). Three assembly schemes were implemented, assemblies per sample, co-assemblies of samples from the same tributary and a unique co-assembly built from all samples. Comparison of the 3 schemes lead to the global co-assembly being chosen for the rest of the analysis. Contigs smaller than 500 nucleotides were discarded. Gene prediction was carried out using prodigal v2.6.3, with option - p meta (Hyatt et al, 2015). A selection of 36 Single copy Core Genes (SCG) were annotated through rpsblast v2.9.0 using pssm formatted COG database made available by the CDD (Tatusoy et al 2000; Lu et al, 2020). Best hits were filtered so that hit span at least 50% of the query with an e-value smaller than 1e-5. Additionally, ARG and *intI1* annotation was completed using diamond v0.9.9 (Buchfink et al, 2015). Databases for both were sourced from (Lee et al, 2020), clustered respectively from of CARD (Jia et al, 2017) and NCBI sequence search. Best hits were filtered using 80% query coverage, 80% percentage identity and 20% subject coverage. Taxonomic annotation was carried out using diamond blastp on a NR database reduced to prokaryotes (Pruitt et al, 2005).

Assembly was converted to contig graph using megahit toolkit, it was then translated to orf graph using result from gene prediction in custom python script. Visualisation of annotations on orf graph is done through bandage v0.8.1 (Wick et al, 2015).

Samples were mapped to contigs using bwa-mem v0.7.17-r1188 (Li, 2013). Contig and genes mean coverage were computed through both samtools v1.5 (Li et al, 2009) and bedtools v2.28.0 (Quinlan et al, 2013). Coverage of functional feature, such as SCG, AMR or *intI1* were obtained by summation of coverage of genes sharing the same annotation. All coverages are normalised by sample, using the median of the 36 SCG coverages.

### Chemical Analysis

For each site 100 mL of each water sample was acidified to pH 3, internal standards were added and the sample was extracted with a solid phase extraction (SPE) vacuum manifold system using Oasis HLB (200mg) cartridges. Thereafter each sample was evaporated to <10 μL, reconstituted to a final volume of 0.1 mL 5% acetonitrile with 0.1% formic acid and 10 μL of each water sample was analysed with liquid chromatography-tandem mass spectrometry (LC-MS/MS). A detailed description of the two analytical methods used to analyse most of the pharmaceuticals, and the validation of each method is shown in (Holden et al, 2020). However, at the time of sample analysis, contamination in the LC-MS/MS system interfered with the analysis of the six biocides included in the study. Instead, an analytical method developed by (Östman et al, 2017) specifically for biocides, on the LC-MS/MS system, was used to analyse the six biocides.

### Geospatial analyses and meteorological data

All geospatial analyses were performed using ArcGIS v10.4 software (ESRI, Aylesbury, UK). To analyse possible drivers of AMR for each sampling site, an upstream catchment of that location was delineated using the hydrology tools to create a flow accumulation and flow direction layer per site, and subsequently a watershed (upstream catchment), using a 50m grid interval hydrological digital terrain model (the Centre for Ecology and Hydrology Integrated Hydrological Digital Terrain Model (IHDTM)) (Morris & Flavin, 1994). A Network Dataset was created from the Thames river data, using the Ordnance Survey MasterMap Water Network 2016 (Ordnance Survey, 2016) and comprised all layers needed for distance measurements, including WWTPs. Data on WWTPs was obtained from the environment agency (Environment Agency, 2012; Environment Agency, 2013). To analyse upstream land use, and more specifically to calculate the percentage land cover of each sample site watershed, the Centre for Ecology and Hydrology Land Cover Map 2015 (Rowland et al, 2017) and Land Cover Plus: Crops Map 2016 were used (NERC CEH, 2016).

Land-cover profiles were calculated for each sampling site based on the identified watershed, comprising the percentage cover for 13 broad habitat classes (Arable horticulture, Acid Grassland, Broadleaf woodland, Calcareous grassland, Coniferous woodland, Freshwater, Heather, Heather grassland, Improved grassland, Inland rock, Neutral grassland, Suburban, Urban). The “Arable horticulture” class was further sub-divided into 12 primary crop categories, and three further major land-cover classes were also calculated by summing over broad habitats (All Grassland (sum of 5 broad habitats), All Woodland (sum of 2 broad habitats), Urban-Suburban (sum of 2 broad habitats)).

For each WWTP in each watershed, data on the Population Equivalent (overall sewage load), resident and non-resident populations and industry load (all in terms of the people producing the equivalent sewage load assuming 200l of sewage flow containing 60g of BOD per person per day) were tabulated together with the river distance (m) from the WWTP to the sampling site, the dry weather flow at the WWTP (m^3^ per day) and the general classification of sewage treatment plants (WWTP Type: SA - Secondary Activated Sludge; SB - secondary biological filter; TA - Activated Sludge with tertiary treatment; TB - Biological filter with tertiary treatment). Simple summaries of these data were calculated for each watershed (sampling site), including total numbers of WWTPs and number for each Type, mean, minimum and maximum distance, and mean and maximum PE. Based on the modelling approach developed in (Amos et al, 2015), summaries of the potential aggregated impact of upstream WWTPs were also calculated – these weighted the impact of the load (PE) from each WWTP by the inverse of the distance raised to a power (0, 0.35 (close to the value estimated in the model fitting in (Amos et al, 2015), 0.5 or 1), allowing for different decay functions (with a power of zero indicating no impact of distance). Where (Amos et al, 2015) only considered WWTPs within a 10km radius of the sampling site, summaries of the numbers of WWTPs and the inverse distance-weighted aggregated PE were calculated for WWTPs within 10km, 20km, 30km, 40km and 50km upstream as well as for the whole upstream watershed. Data were also tabulated for each sampling site (watershed) for the number and density of septic tanks, and for the number and mean upstream distance for fish farms.

### Statistical analysis and model development

Initial analysis of chemical data used a mixed-effects model with Tukey’s multiple comparison tests to assess differences in trimethoprim (TMP) and sulfamethoxazole (SMX) concentrations between seasons (summer & winter) and between twenty sample sites (Coln1, Coln2, Coln3, Coln4, Leach1, Leach2, Key1, Ray1, Ray6, Ray4, Ray3, Thames1, Thames3, Thame7, Thame2, Cut1, Cut2, Cut4, Cut6, Thames6).

Analysis of geospatial data relating to the distance (metres) of the twenty sample sites downstream from waste-water treatment plants (WWTPs) and the concentration of TMP was carried out using Spearman’s rank correlation coefficient due to the non-parametric nature of the data. Both variables were log-transformed prior to analysis. Analysis of the difference in population equivalence (PE) between sites with the highest and lowest TMP levels was carried out using an unpaired t-test with Welch’s correction.

Analysis of metagenomic data used multiple t-tests with Bonferri-Dunn correction to assess differences in ARG (*sul1*, *sul2*, *dfrB1/6*, *dfrB2/3*, *dfrE*) and *intI1* prevalence between seasons (summer and winter). Two-way ANOVA was also applied to assess differences in the prevalence of aforementioned ARGs & *intI1* between seasons and sites with the four highest (Cut4, Ray4, Cut6, Ray6) and four lowest (Coln1, Coln2, Coln3, Leach1) TMP concentrations. All analysis was performed using GraphPad^®^ Prism version 8 and R^®^ software. For all tests, p values of statistical significance are signified by * = p<0.05, ** = p<0.005, *** = p<0.0005 & **** p<0.0001.

### Maps

Maps shown in Figure 4 were created using ArcGIS v10.4 software. Geographic coordinates of sites were taken from (Ordnance Survey, 2016).

## Results

### Analysis of TMP and SMX levels

The levels of TMP and SMX identified in the water at the twenty different sites across Thames catchment differed greatly. Furthermore, there was also a significant seasonal variation in the levels of TMP and SMX identified at individual sites (Figure 2).

**Figure 2.**
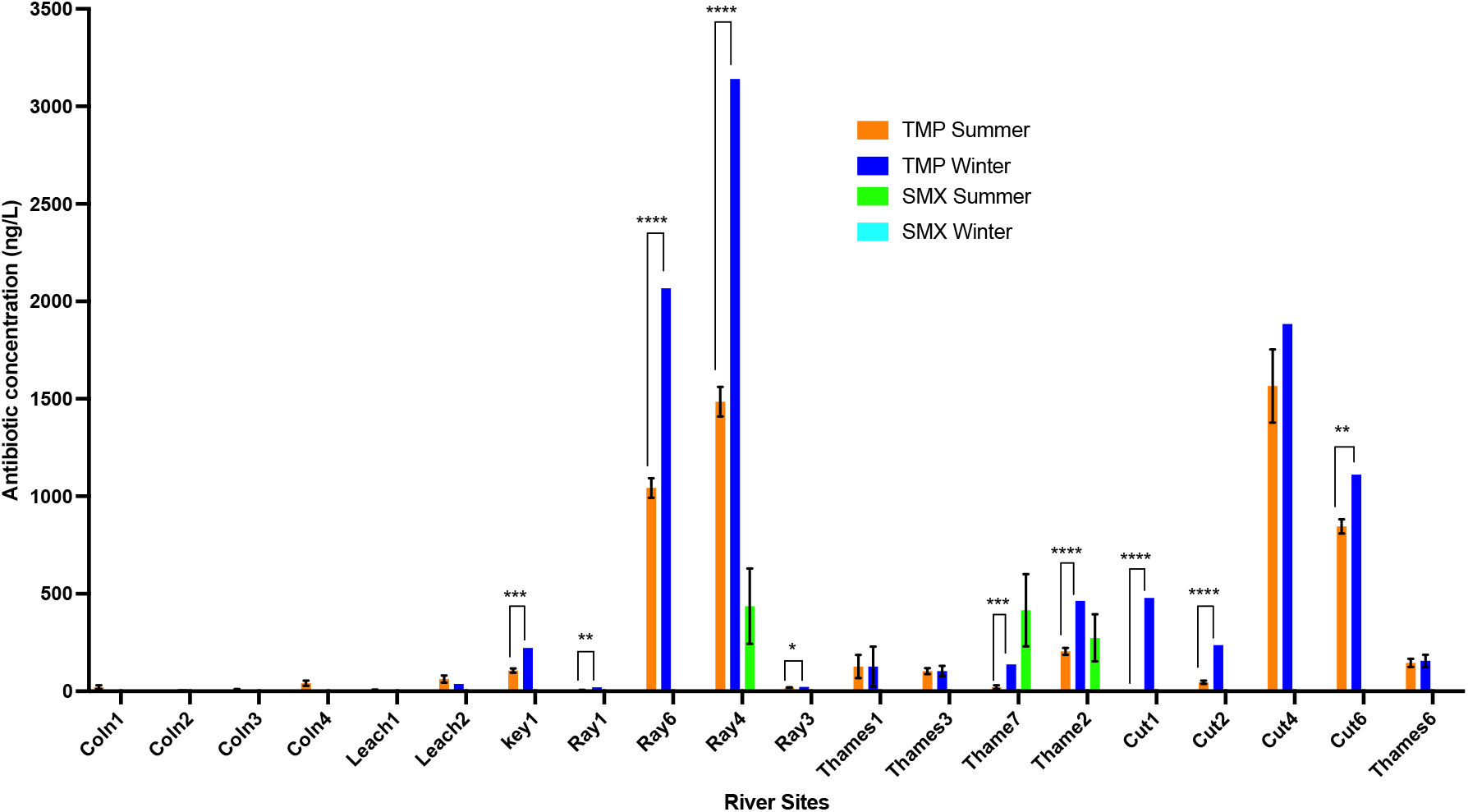
The concentrations of TMP and SMX at twenty sites across the Thames catchment in summer (n=6) and winter (n=3) (SEM). The sites were ordered by geographical location across the catchment, with Coln1 being furthest East and Thames6 being furthest West.

**Figure 3.**
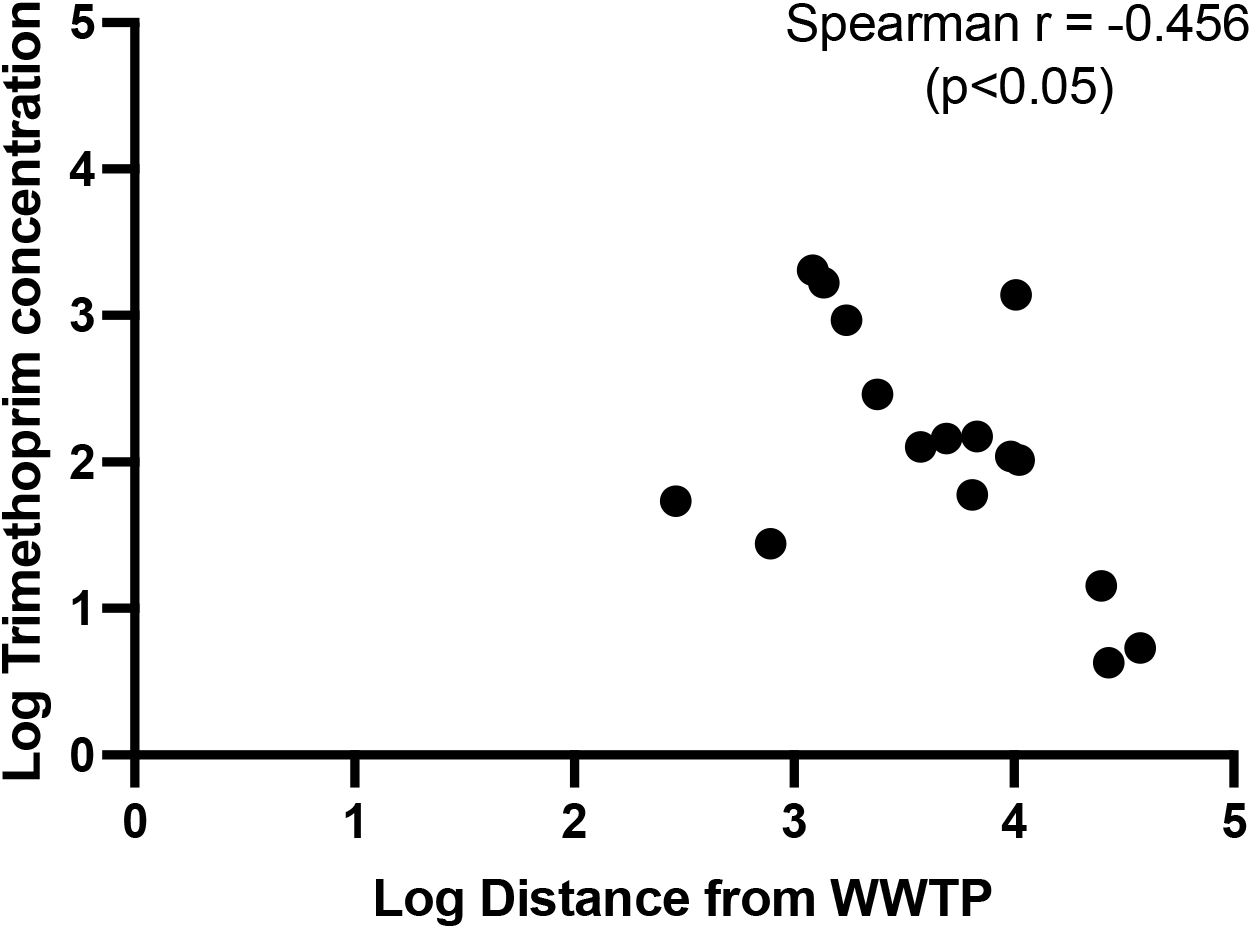
Relationship between distance of sample sites downstream from a WWTP and the concentration of TMP across 20 sites on the Thames catchment.

**Figure 4.**
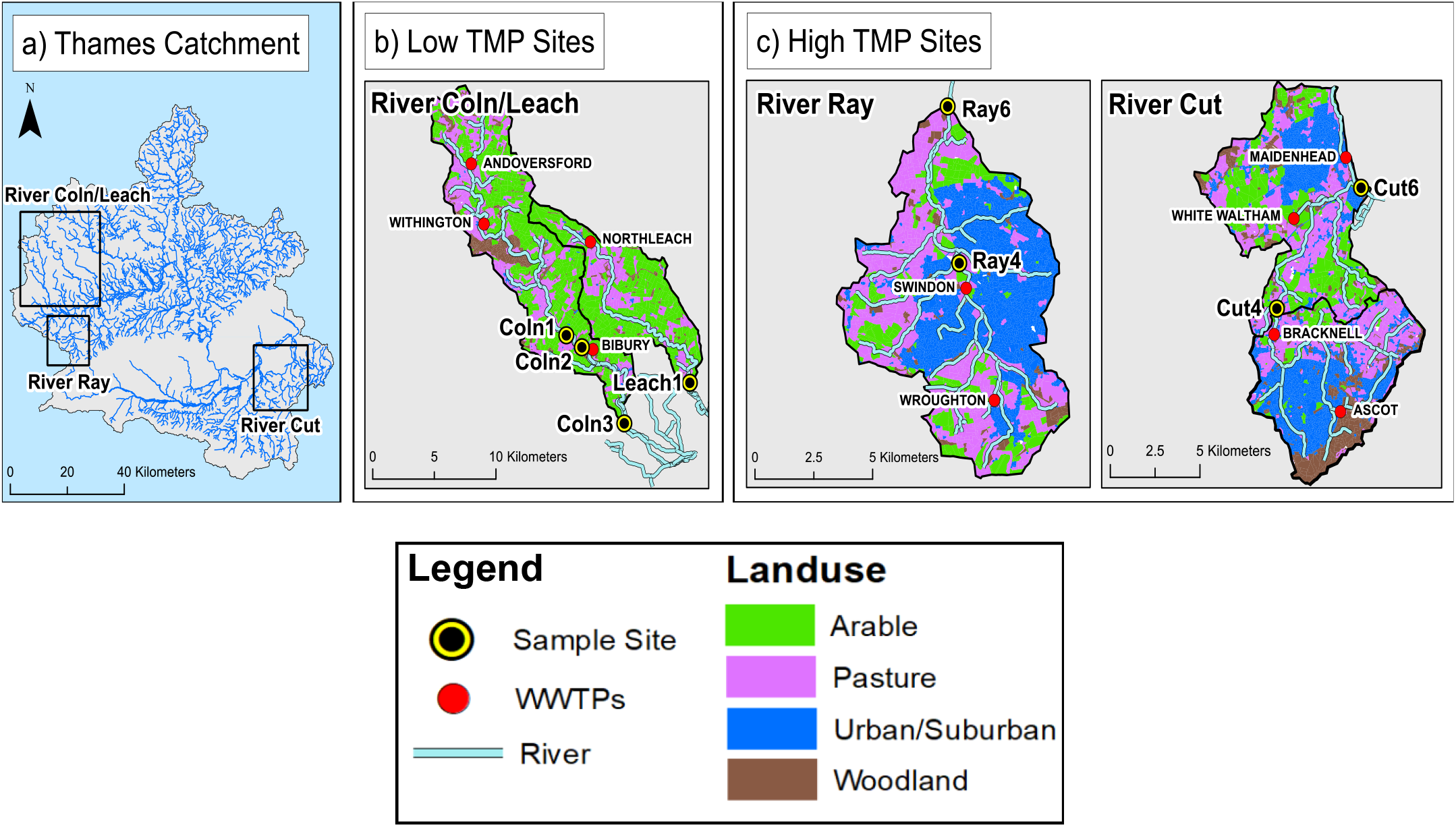
A) Locations of the Rivers Cut, Ray, Coln and Leach on the Thames catchment.
B) Locations and land-use corresponding to the four sites with the lowest TMP concentrations.
C) Locations and land-use corresponding to the four sites with the highest TMP concentrations.

TMP concentrations were higher than SMX concentrations at all sites other than Thame7. Ten of the twenty sites had significantly higher TMP in winter compared to summer (mixed effects model). Four sites had levels of TMP exceeding 750ng/L (Ray4, Ray6, Cut4 and Cut6), with Ray4 providing a TMP concentration of 3000ng/L. Aside from Cut4, the level of TMP was significantly higher in winter compared to summer for these sites (mixed effects model).

For three sites (Ray4, Thame7 and Thame2), the concentration of SMX was higher in summer compared to winter, but these differences were not significant (mixed effects model).

A significant negative correlation was observed between the distance of sites downstream from the nearest WWTP and the TMP concentration across 20 sites on the Thames catchment (Spearman r = −0.456 & p<0.05). This relationship was further explored to determine where the highest and lowest TMP levels were detected (Table 1).

<u>**Table 1**</u>- A sub-set of four sites that showed the highest and lowest TMP concentrations from the 20 sites. Of this sub-set, only one of the sites showed a recordable concentration of SMX (Ray4), so this data was not shown in the table. TMP concentrations shown were averaged across summer/winter seasons.

#### * Secondary

‘Treatment methods include biological filtration (including conventional filtration, rotating biological contactors and root zone treatment (where used as a secondary treatment stage), alternating double filtration and high rate filtration’. (Sewage Treatment in the UK, 2020).

#### * Tertiary

**’**Works with a secondary activated sludge process whose treatment methods also include rapid-gravity sand filters, moving bed filters, pressure filters, nutrient control using physico-chemical and biological methods, disinfection, hard COD and colour removal, where used as a tertiary treatment stage’ (Sewage Treatment in the UK, 2020).

| River Site | Nearest upstream WWTP | Distance from WWTP (m) | WWTP Treatment Type* | Human population served | Trimethoprim concentration (ng/L) |
| --- | --- | --- | --- | --- | --- |
| Cut4 | Bracknell | 1361 | Tertiary | 90,000 | 1671 |
| Cut6 | Maidenhead | 1540 | Tertiary | 87,400 | 934 |
| Ray4 | Swindon | 1213 | Tertiary | 210,000 | 2037 |
| Ray6 | Swindon | 10208 | Tertiary | 210,000 | 1384 |
| Coln1 | Withington | 16677 | Secondary | 230 | 14 |
| Coln2 | Withington | 18688 | Secondary | 230 | 4 |
| Coln3 | Bibury | 9479 | Secondary | 600 | 5 |
| Leach1 | Northleach | 20270 | Secondary | 1,670 | 4 |

Three of the four sites with the highest TMP concentrations were located within 2000 metres of WWTPs (Table 1) as expected from the data in Figure 4. The four sites with the highest TMP concentrations were located closest to tertiary WWTPs, whereas the all of the four sites with the lowest TMP concentrations were located closest to secondary WWTPs. Two of the four sites with the highest TMP concentrations (Ray4 and Ray6) were located closest to the same Swindon WWTP.

### Analysis of land use and population equivalence (PE)

Land use and PE data were assembled for the four sites with the highest and lowest TMP concentrations across the Thames catchment, as shown in Figures 4, 5 and 6.

**Figure 5.**
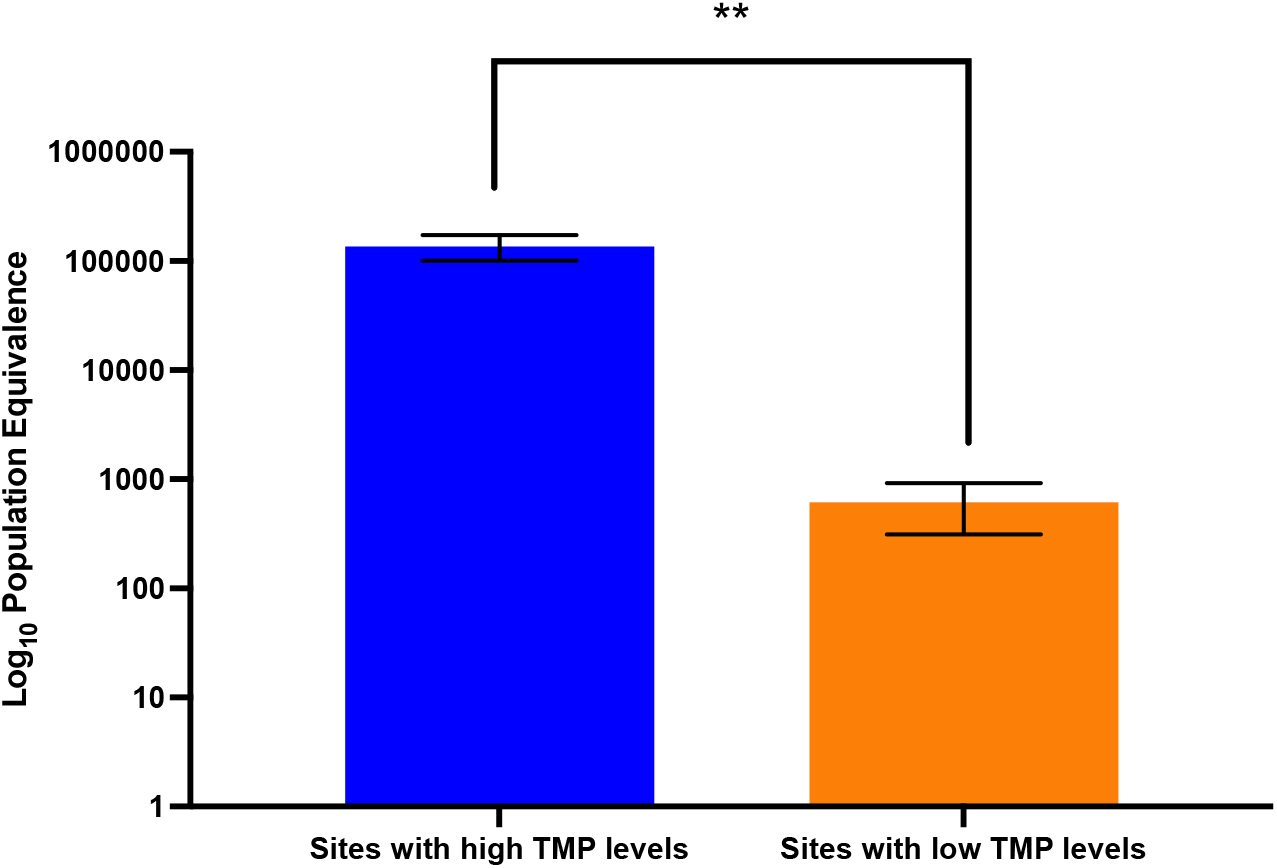
The average Log_10_ population equivalence (PE) for four sites with the highest TMP concentrations (left) and four sites with the lowest TMP concentrations (right) (n=4) (SEM).

**Figure 6.**
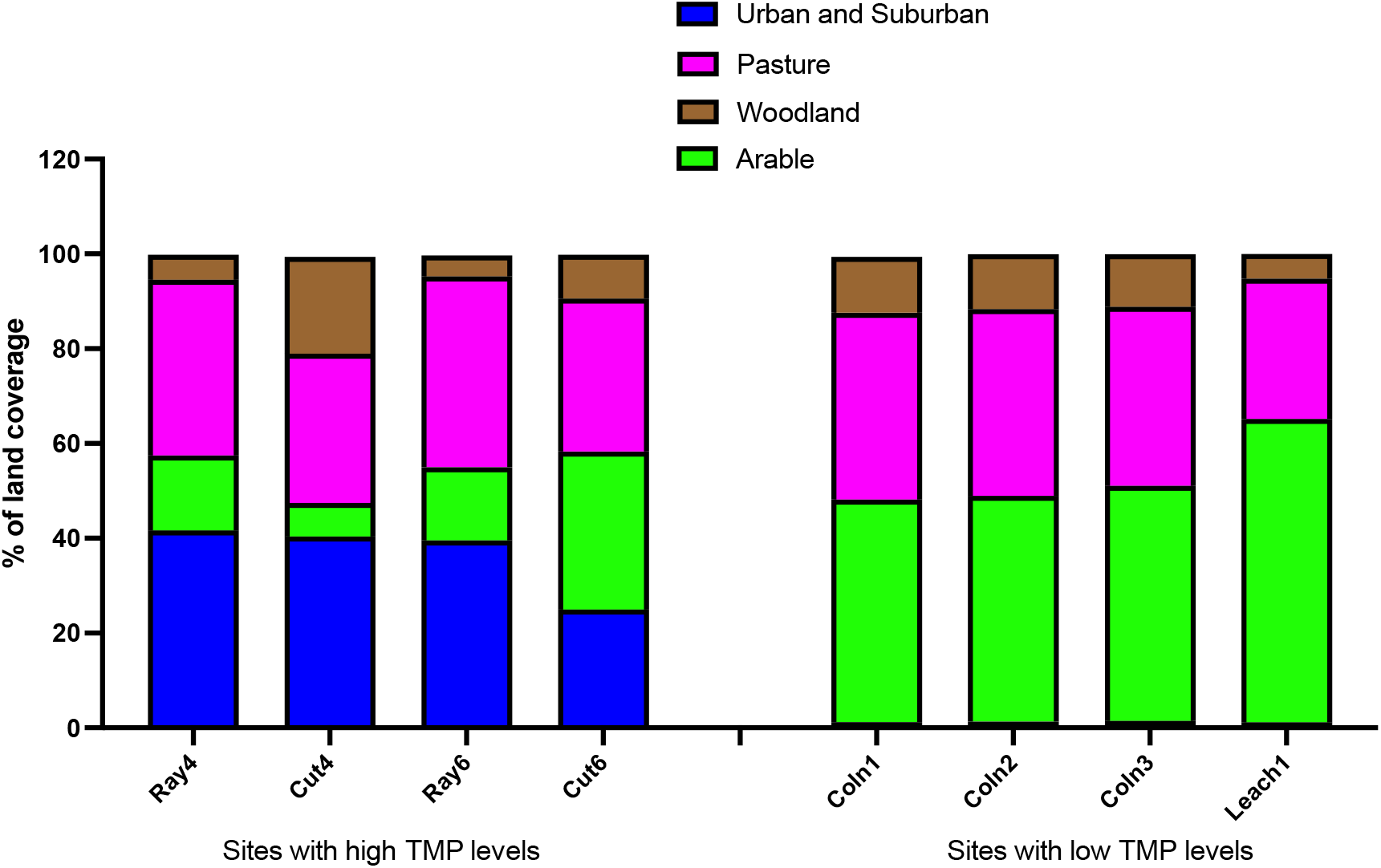
Percentage of land coverage within the catchment of four sites with the highest TMP levels (left) and the lowest TMP levels (right).

Examination of the land-use data in proximity to sites with highest and lowest TMP levels indicate urbanization correlating with tertiary WWTPs and high TMP levels (Figure 4). The data also shows that the proportion of arable and pasture-based land coverage was higher within the catchment of sites furthest from WWTPs.

The average PE for the four sites with the highest TMP concentrations was significantly higher than the PE for the four sites with the lowest TMP concentrations (unpaired t-test).

The proportion of urban/suburban land coverage was found to be higher in the catchment of the four sites with the highest TMP concentrations compared to the four with the lowest TMP concentrations. By contrast, the proportion of arable land coverage was higher across the catchment of sites with the lowest TMP concentrations compared to sites with the highest TMP concentrations.

### Prevalence of ARGs

Only two SMX ARGs and three TMP ARGs were found at detectable levels in the metagenomes derived from river sediment (Figure 7) and of these *sul1* was detected at significantly higher levels in the winter. These were compared with *intI1* (CL1 marker gene) prevalence (Figure 7).

**Figure 7.**
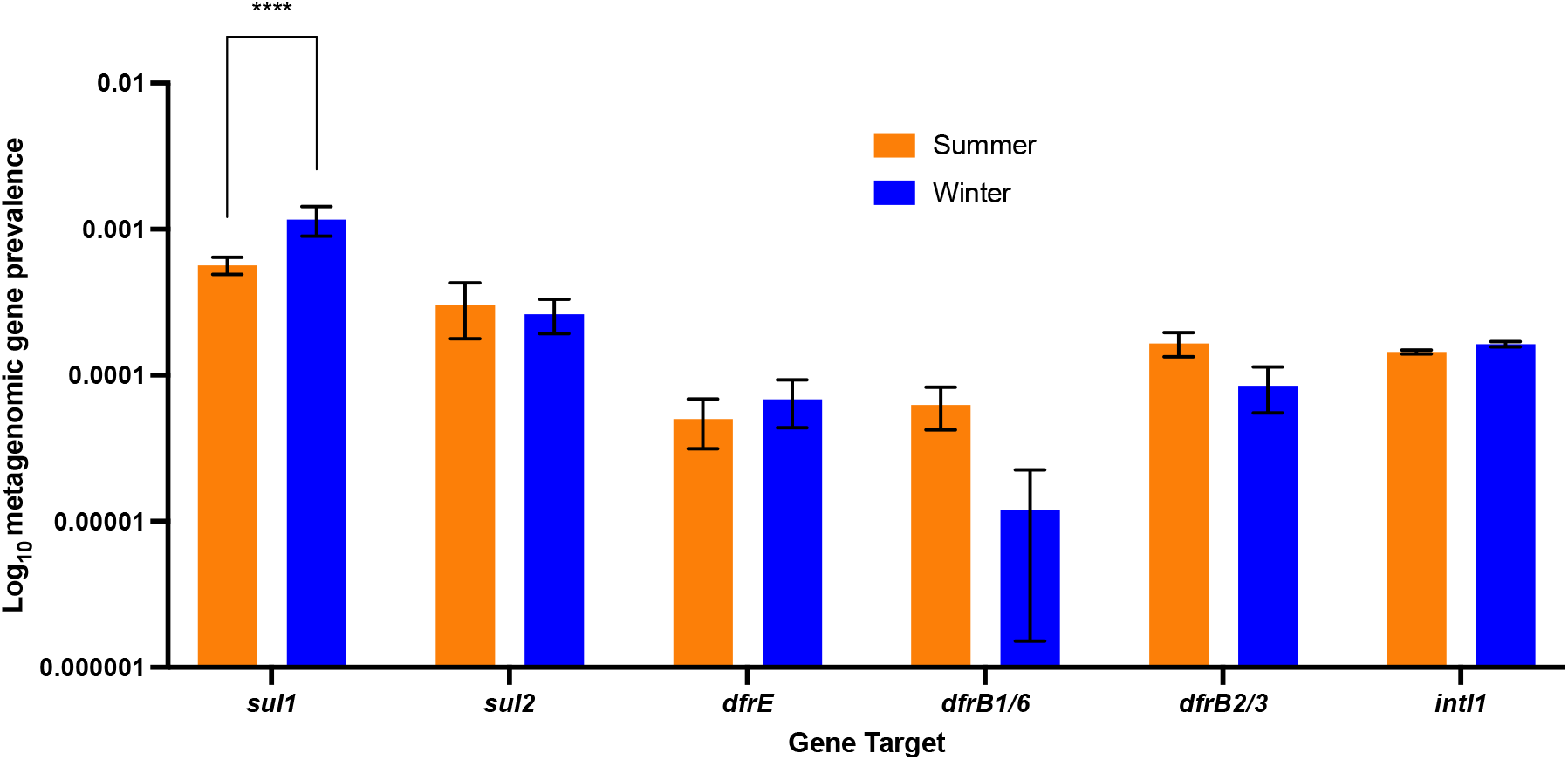
Prevalence of ARGs relevant to TMP/SMX resistance across 20 sites on the Thames catchment in summer (n=6) and winter (n=3) (SEM).

**Figure 8.**
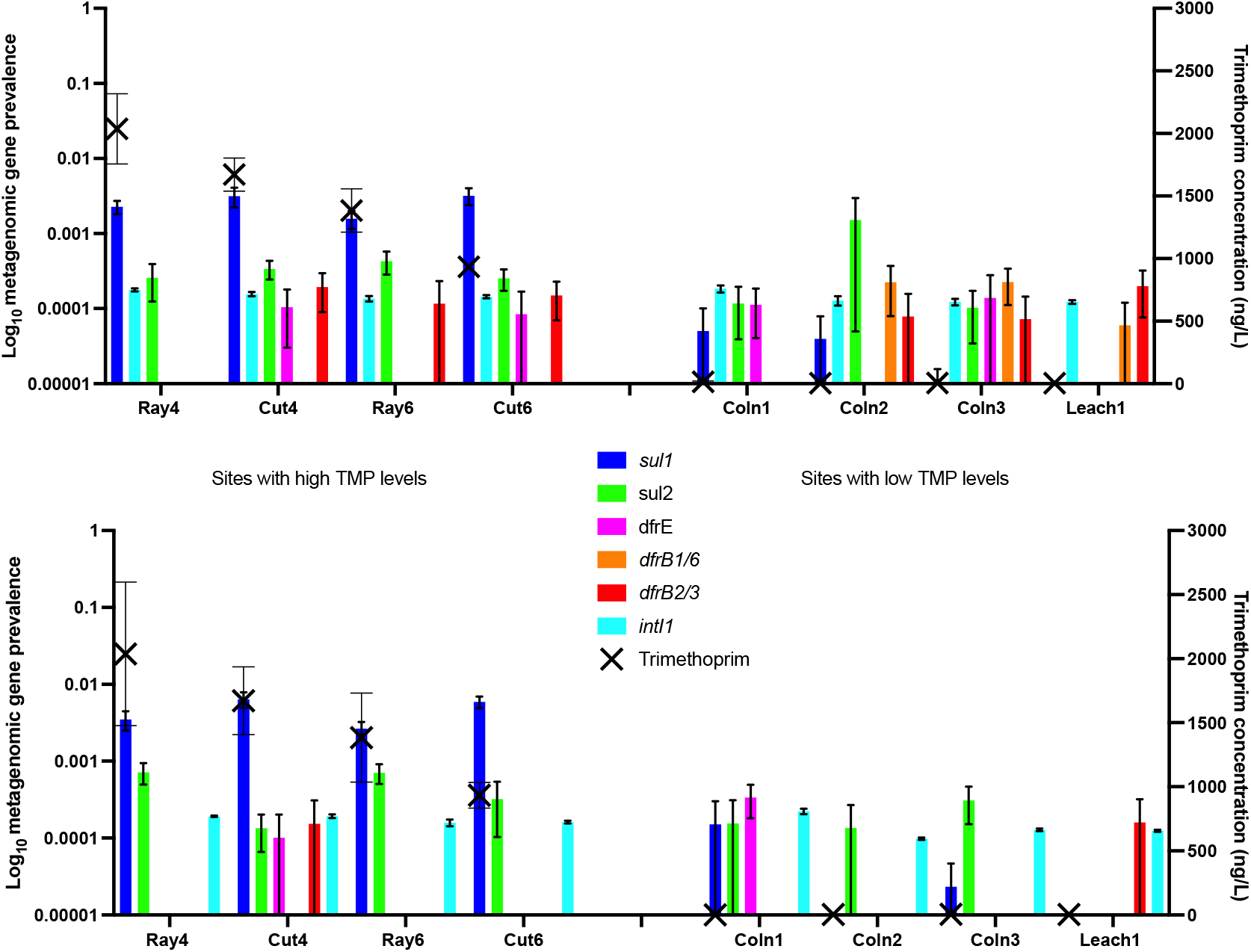
Prevalence of ARGs relevant to TMP/SMX and *intI1* at sites with the four highest and four lowest TMP levels in summer (n=6) (above) compared to winter (n=3) (SEM).

The *sul1* gene showed the highest overall prevalence compared to the four other ARGs relevant to TMP and SMX and *intI1*. *sul1* was also the only gene to show significant variation between the summer and winter seasons (multiple t-tests).

The two ARGs relevant to SMX and the three ARGs relevant to TMP were then analysed in comparison with the levels of TMP antibiotic identified at the sites with the four highest and four lowest TMP levels (same sub-set as Table 1). These were compared with *intI1* (CL1 marker gene) prevalence.

The prevalence of *sul1* was higher at sites Ray4, Ray6 and Cut4 in winter compared to summer. At these four sites, the prevalence of *sul1* was higher than any of the prevalence of any other genes in both winter and summer. Coln2 and Coln3 showed significantly higher prevalence’s of *dfrB1/6* in summer compared to winter (two-way ANOVA). The prevalence of *sul2* at Coln2 was significantly higher in summer compared to winter (two-way ANOVA).

## Discussion

Over the last two decades, the occurrence of antibiotics in water bodies and the potential for the development of antimicrobial resistance (AMR) as a result has become an issue of concern (Khan et al, 2013). Particularly in urban areas, waste-water treatment plants (WWTPs) are among the main recipients of antibiotics, some of which will persist after treatment and can be released into nearby rivers (Rodriguez-Mozaz et al, 2020).

The levels of two clinically relevant antibiotics (trimethoprim (TMP) and sulfamethoxazole (SMX)) were analysed at river sites during this study. Results showed that nineteen of the twenty river sites analysed showed higher levels of TMP compared to SMX. TMP and SMX can be co-prescribed as co-trimoxazole, which might suggest that their levels identified in rivers might be similar. However, clinical antibiotic surveillance data shows that TMP is more commonly prescribed as a single dose antibiotic (Ashiru-Oredope et al, 2014). By contrast, veterinary surveillance data suggests that TMP is co-prescribed with ‘sulfonamides’ in high levels in agricultural settings (Bos et al, 2019). As a result, it is possible that TMP levels were higher than SMX levels at these sites because the two drugs are more often being used for different purposes and therefore more likely to be found at different river sites.

Another possible reason why TMP was significantly higher than SMX at certain sites could be the differing biodegradation profiles that the drugs display in the environment (Thiebault, 2020). One study used batch reactors containing wastewater matrices to show that the biodegradation rate of SMX was significantly higher than TMP over a 21-day period (Pérez et al, 2005). Most WWTPs are designed to extract nutrients and easily degradable carbon compounds. However, it has been shown that they have limited capacities to remove pharmaceuticals such as TMP, making them sources of TMP in rivers and streams (Metcalfe et al, 2010; Sengupta et al, 2014). The presence of antibiotics in the treatment process can inhibit the biological activity of activated sludge, meaning that the antibiotics themselves are not degraded effectively (Al-Riyami et al, 2018). As a result, it is possible that SMX is more readily degraded than TMP as part of the WWTP process and therefore was found in higher concentrations at certain sites.

As a result of the incapacity of conventional treatment systems to remove emerging contaminants such as antibiotics during treatment, studies have indicated that urban-associated river sites often have the highest concentrations of antibiotics (Chen et al, 2018; Lundborg et al, 2017). This would explain why TMP and SMX were both detected at river sites in this study, particularly as those sites with the highest TMP levels all had a high level of urban/suburban land coverage associated with them. The significant negative correlation between levels of TMP and the distance of a site from a WWTP further suggests that the TMP being identified in rivers coming from WWTPs. Furthermore, in areas of high human population, it has been shown that excretion by humans contributes to most of the pharmaceutical waste that ends up in domestic sewers and is eventually transported to WWTPs (Tehrani and Gilbrade, 2018). This would explain the fact that the PE was significantly higher at the four sites with the highest TMP concentrations compared to those with the lowest TMP concentrations.

During this study, four river sites (Cut4, Cut6, Ray4 and Ray6) were shown to have elevated levels of TMP. Despite showing significantly higher levels of TMP compared to all other sites, they did not show a significantly higher prevalence of the ARGs that are known to confer resistance to TMP (*Dfr* genes). *DfrB1/6* for example, was only detected at sites with the lowest TMP levels. This suggests that the concentrations of TMP at river sites are not directly selecting for resistance, which contrasts with previous reports which have shown that sub-Minimum inhibitory concentrations (MIC) of antimicrobials can select for resistance in environmental settings (Ter Kuile et al, 2016; Winstrand-Yuen et al, 2018; Opatowski et al, 2010). Interestingly, sites with the highest levels of the SMX associated ARG *sul1* did not show high levels of SMX but did show high levels of TMP. One possible explanation for why this happens is that TMP is indirectly selecting for *sul1* by selecting for the presence of class 1 integrons (CL1s), as CL1s are known to carry the *sul1* gene. CL1s which have been shown to confer resistance to several antibiotics, including SMX, TMP, amoxicillin (AMX), tetracycline (TCN) and erythromycin (ERY) (Langata et al, 2019; Ndi and Barton, 2011; Callejas et al, 2019). Our previous data showed no significant correlations between TCN and ERY concentrations with *sul1* prevalence, but we do not have chemical data available relating to AMX (Holden et al, 2020). It is also possible that concentrations of AMX are present at these river sites and they are also able to select for CL1s and therefore *sul1*. This effect has been noted in a previous study (Ironmonger et al, 2018). However, based solely on the data shown in this study, given that the prevalence of CL1s did not differ significantly between river sites or seasons like *sul1* did, it seems unlikely that this is the case. Another possible explanation is that both TMP and *sul1* are released to rivers following the waste-water treatment process at the same time, and therefore were both detected by coincidence. This would mean that the relationship between TMP and *sul1* noted in this paper and our previous study is purely correlative (Holden et al, 2020).

Data from this study showed seasonal variation in the prevalence of individual ARGs and the concentrations of TMP/SMX antibiotics at river sites. For example, the prevalence of *sul1* over the 20 sites was significantly higher in winter compared to summer. Also, the concentration of TMP was significantly higher in winter compared to summer for ten of the twenty sites analysed. Increased prescription/use of TMP in the population during the winter months could increase the concentrations detected at river sites and potentially the prevalence of ARGs. A recent study indicated that prescription rates for antibiotics (including TMP) by General Practitioners (GPs) in the West Midlands UK increase consistently in winter compared to the three other seasons (Ironmonger et al, 2018). An increase in antibiotic consumption can increase the antibiotic concentration found in sewage, which will eventually be treated by WWTPs (Pärnänen et al, 2019). If antibiotics are not successfully removed during sewage treatment processes, then the higher concentrations of TMP shown in winter compared to summer could be due to increased antibiotic usage/consumption. However, with regards the chemical analysis of TMP and SMX, it must be acknowledged that the freshwater samples used were collected over two summer seasons (summer 2015 and summer 2016) and only one winter season (winter 2016). Also, the concentrations of chemicals determined at each site are only reflective of the concentration of that chemical at one particular time, with no corrections made for the flow rate of the individual river site or data on sewage spills from WWTPs. As a result, the chemical data shown is potentially not reflective of the whole seasons analysed and the differences between TMP concentrations in summer and winter is potentially more of a temporal effect as opposed to a seasonal one.

One site in this study; Coln2 showed a showed significantly increased prevalence of *sul2* in summer despite having minimal concentrations of TMP and SMX, which contrasted with its low *sul1* prevalence. *Sul2* is a sulfonamide resistant dihydropteroate synthase of gram-negative bacteria, generally found on small plasmids, whereas *sul1* is generally found linked to other resistance genes on CL1s (Antunes et al, 2005; Jia et al, 2017). According to the National Office of Animal Health (NOAH), there are two different sulfonamide classes in addition to SMX that can be prescribed in combination with TMP to treat infections in farm animals: sulfadiazine (SDZ) and sulfadoxine (SX) (NOAH, 2020). TMP-SDZ is a short-acting bacteriostatic antibiotic that is typically applied to treat infections in horses (Ensink et al, 2003). TMP-SX is a bacteriostatic antibiotic typically used to treat respiratory tract infections in cattle (Dunkley, 1994). Previous studies have noted increased abundance of the *sul2* in agricultural samples containing SDZ (Jechalke et al, 2012; Kopmann et al, 2012). As a result, is possible that the increased prevalence of *sul2* shown at Coln2 in spite of the low SMX and TMP concentrations is driven by an agriculturally relevant sulfonamide such as SDZ, which was not targeted by the chemical analysis of this study. This is further emphasised by the fact that there was no relationship between *sul2* and *intI1* prevalence in this study, as this shows that *sul2* genes are not emerging on CL1s alongside clinically relevant ARGs. However, it must be acknowledged that the available UK antibiotic surveillance data for veterinary applications only shows data for ‘sulfonamides’ as a general class, as opposed to distinguishing between individual drugs such as SMX and SDZ (Bos et al, 2019). As a result, it is difficult to determine the actual levels of SDZ that are being applied in agriculture. Also, despite the isolated high levels of *sul2* at Coln2, this study showed that across all 20 sites on the Thames catchment, *sul1* was significantly more prevalent than *sul2.*

## Conclusions

This study involved comparative analysis of large metagenomic, chemical and geospatial datasets to investigate the impact of antimicrobials such as trimethoprim (TMP) on the profile of antimicrobial resistance and the wider river microbiome. The following conclusions were drawn:

1) Nineteen of the twenty sites analysed showed higher levels of trimethoprim (TMP) than sulfamethoxazole (SMX), except for Thame7.
2) There was a significant negative correlation between the distance of river sites from waste-water treatment plants (WWTP) and the levels of TMP detected.
3) Of the twenty river sites analysed, four showed particularly high levels of TMP. For three of these four sites, TMP levels were significantly higher in winter compared to summer.
4) *DfrE*, *DfrB1/6*, *DfrB2/3 and intI1* prevalence at the four sites with the highest TMP levels were not significantly higher than those with the lowest TMP levels. *DfrE*, *DfrB1/6*, *DfrB2/3* was significantly higher in summer compared to winter. *intI1* prevalence showed no significant seasonal variation.
5) The four sites with high TMP levels showed significantly higher PE (population equivalence) levels than the four sites with the lowest TMP levels.
6) The land coverage of the four sites with the highest TMP levels were significantly more urban/suburban than the four sites with the lowest TMP levels. By contrast, the land coverage of the four sites with the lowest TMP levels was significantly more arable than those with the highest TMP levels.
7) *sul1* prevalence at the four sites with the highest TMP levels were significantly higher than those with the lowest TMP levels. *sul1* prevalence was also significantly higher in winter compared to summer.
8) *sul2* prevalence at one of the sites with the lowest TMP levels (Coln2) was significantly higher than those with the highest TMP levels. *sul2* prevalence at the Coln2 was also significantly higher in summer compared to winter.

### Future Work

Our previous work has shown evidence of a significant positive correlation between TMP levels and the prevalence of *sul1* at sites across the Thames catchment (Holden et al, 2020). Data from this study pinpointed the sites with the highest levels of TMP and the highest prevalence of *sul1.*

We have also shown that sites with the highest TMP concentrations and highest *sul1* prevalence are generally located closer to WWTPs than those with low TMP concentrations. Furthermore, sites with the highest TMP concentrations have a higher PE, are more urban/suburban and that their TMP concentration/*sul1* prevalence is higher in winter compared to summer. Despite this, there is still no previously explained mechanism as to why levels of TMP would select for *sul1* directly.

definitive evidence of a causative impact of TMP concentrations on the prevalence of ARGs such as *sul1*. Future work will aim to look at whether or not the concentrations of TMP and SMX recorded in this study can have a significant causative impact on the prevalence of relevant ARGs.

## Notes

### Competing Interest Statement

The authors have declared no competing interest.

